# Modelling Gene-Protein-Reaction Associations on an FPGA

**DOI:** 10.1101/658195

**Authors:** Macauley Coggins

**Keywords:** FPGA, Genome-scale metabolic models, GPRA, Genes, Reactions, FBA

## Abstract

Genome-Scale metabolic models have proven to be incredibly useful. Allowing researchers to model cellular functionality based upon gene expression. However as the number of genes and reactions increases it can become computationally demanding. The first step in genome-scale metabolic modelling is to model the relationship between genes and reactions in the form of Gene-Protein-Reaction Associations (GPRA). In this research we have developed a way to model GPRAs on an Altera Cyclone II FPGA using Quartus II programmable logic device design software and the VHDL hardware description language. The model consisting of 7 genes and 7 reactions was implemented using 7 combinational functions and 14 I/O pins. This model will be the first step towards creating a full genome scale metabolic model on FPGA devices which we will be fully investigating in future studies.

## 2 Introduction

Genome scale metabolic models have gone a long way since the first genomescale metabolic model which was developed in 1995 for Haemophilus influenzae a single celled organism [1]. This model was based on assembly of unselected pieces of DNA from the whole chromosome had been applied to obtain the complete nucleotide sequence. However the preferred approach today is Flux Balance Analysis (FBA). FBA can be traced back to Papoutsakis et al who in 1984 demonstrated the first use of flux balance equations from metabolic maps [6]. This approach has an advantage over more traditional approaches as it requires very little information about enzyme kinetics or concentration of metabolites in the organism to be described. Instead it assumes that the system will be steady state ie that fluxes are balanced and that the system will always be optimal. The steady state assumption means that the system can be reduced to a set of linear equations which can then be solved using a linear solver such as Gurobi [5], [3]. The first step in constructing a genome scale metabolic model is to describe the relationship between genes and the reactions they encode for. These are called Gene-Protein-Reaction Associations (GPRAs). These are implemented as boolean rules and depending on the reaction these rules can become quite complex. In their simplest form however they model isozymes and enzyme complexes where an isoenyme can be described as

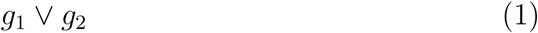

where *g*_1_ and *g*_2_ are the genes coding for the reaction enzyme.

Enzyme complexes can be described as

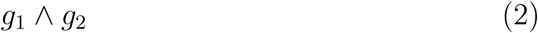

One real world example of a GPRA would be (HGNC:6541 *∨* HGNC:6535 *∨* HGNC:19708) which would describe the reaction Lactate Dehydrogenase. All in-silicos models so far have been software based however dedicated hardware such as FPGAs may be a promising alternative. FPGAs as a platform would provide several advantages as they would provide faster computational speeds. Given that GPRAs are essentially boolean rules these could be modelled naturally on an FPGA using in built logic.

## 3 Materials and Methods

### 3.1 Selection of GPRAs to be modelled

For this research pre-existing GPRAs had to be found from pre-existing literature. GPRA’s were found from Virtual Metabolic Human (VIHM) a site that curates genes, reactions, metabolites and diseases as well as provide information on how all these components interact including GPRAs [4]. For this research GPRAs and reactions that were associated with Lactate Dehydrogenase were selected and are outlined in Table 1. Furthermore Lactate Dehydrogenase is an enzyme that catalyzes the conversion of pyruvate to lactate. The main reason for selecting these are that many genes involved in Lactate Dehydrogenase are also involved in a number of other reactions. Furthermore Lactate Dehydrogenase is a notable enzyme that has been studied for its potential role as a biomarker in tumours [2].

**Table 1:** Modelled Reactions and their GPRAs. Note the gene IDs are HGNC

| Reaction | GPRA |
| --- | --- |
| LDHLM | 6541 or 6535 or 19708 |
| LDHL | (6541 and 6535) or (28335) or (6544) or (6535) or (30866) or (6541) or (21481) |
| 2HBO | (21481) or (6544) or (30866) or (6541 and 6535) or (28335) or (6541) or (6544) |
| GLX01 | (6535) or (30866) or (6541 and 6535) or (19708) or (21481) or (6541) or (28335) or (6544) |
| HPYRR2x | (6544) or (19708) or (6541) or (28335) or (6541 and 6535) or (21481) or (6535) |
| MCLOR | (21481) or (6541 and 6535) or (30866) or (6544) or (6541) or (28335) or (6535) |
| R0173 | (6541) or (6535) or (6544) |

### 3.2 Design and Synsthesis on Quartus II

Quartus II 64-bit is software for analysis and synthesis of HDL designs. Using Quartus II GPRAs were implemented using VHDL a hardware description language used in the synthesis of HDL designs. Genes were implemented as Std logic input signals using while reactions were implemented as Std logic output signals.

GPRAs were then mapped to the reactions using Boolean combinations of genes. Due to the boolean nature of the model gene expression of any gene is simplified to an On/Off state. If a gene signal is true then this corresponds to that particular gene being knocked in. The same can be said for reactions where if the reaction output is true then the reaction is turned on. Next the GPRAs were compiled and a vector waveform file created. A Block Diagram was also created as seen in Figure 1. The vector waveform file provides a way to analyze the functionality of the compiled design whose output is a timing diagram allowing the user to see the inputs and outputs as seen in Figure 1. In the waveform file the genes were grouped together and counted from 0000000 to 1111111 in grey code in order to go through each combination of gene knock-in/knock-out

**Figure 1:**
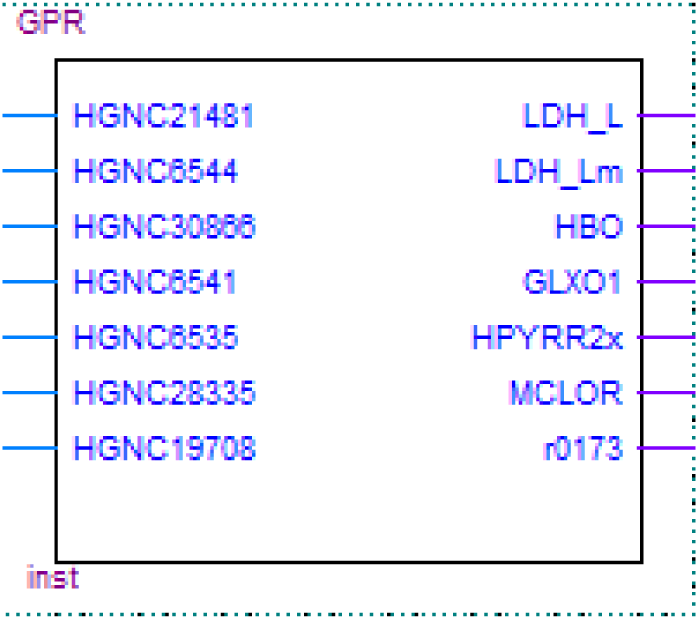
Block Diagram of GPRA Model. Note the genes as input signals to the left and reactions as outputs to the right.

## 4 Results

The results shown in Figure 2 show all combinations of gene knock-in and knock-out were simulated in 989ns with each knock-in/knock-out taking 10ns. The waveform shows that reactions were modelled correctly with no logical hazards. However the risk of hazards is increased as the time between input changes is decreased. Analysis and Synthesis summary shown in Figure 3 shows that the model is very compact as it uses only 7 combinational functions and 14 pins.

**Figure 2:**
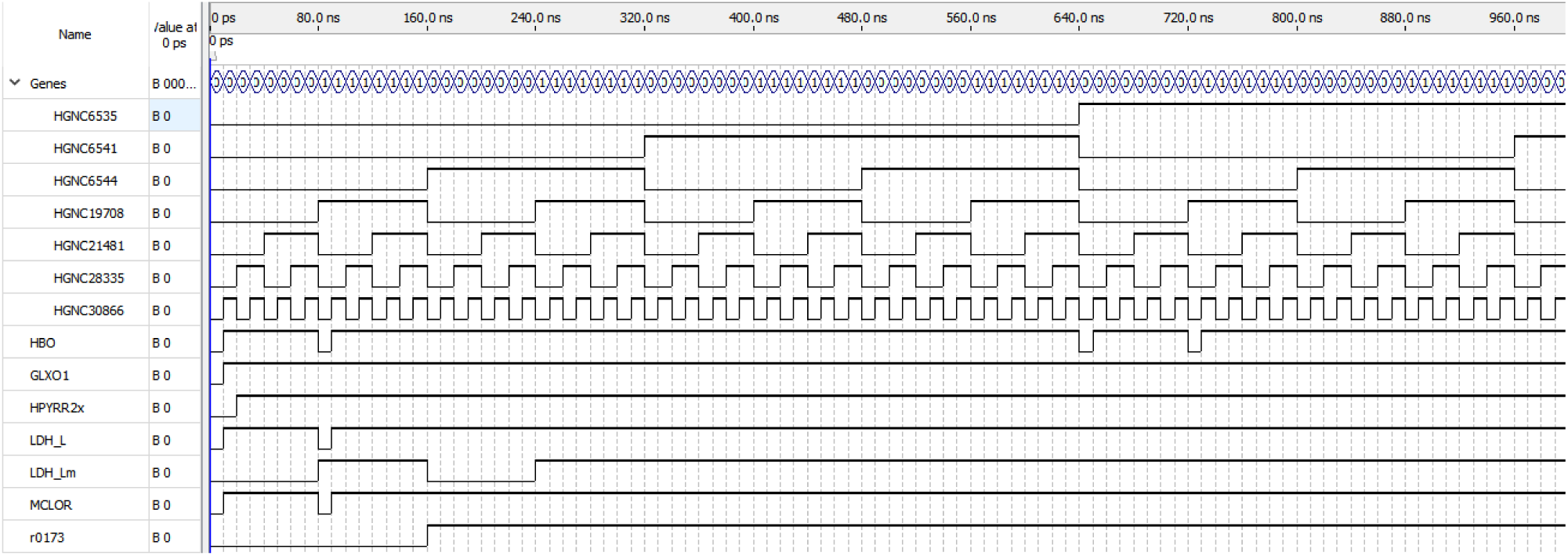
Timing Diagram in Quartus II.

**Figure 3:**
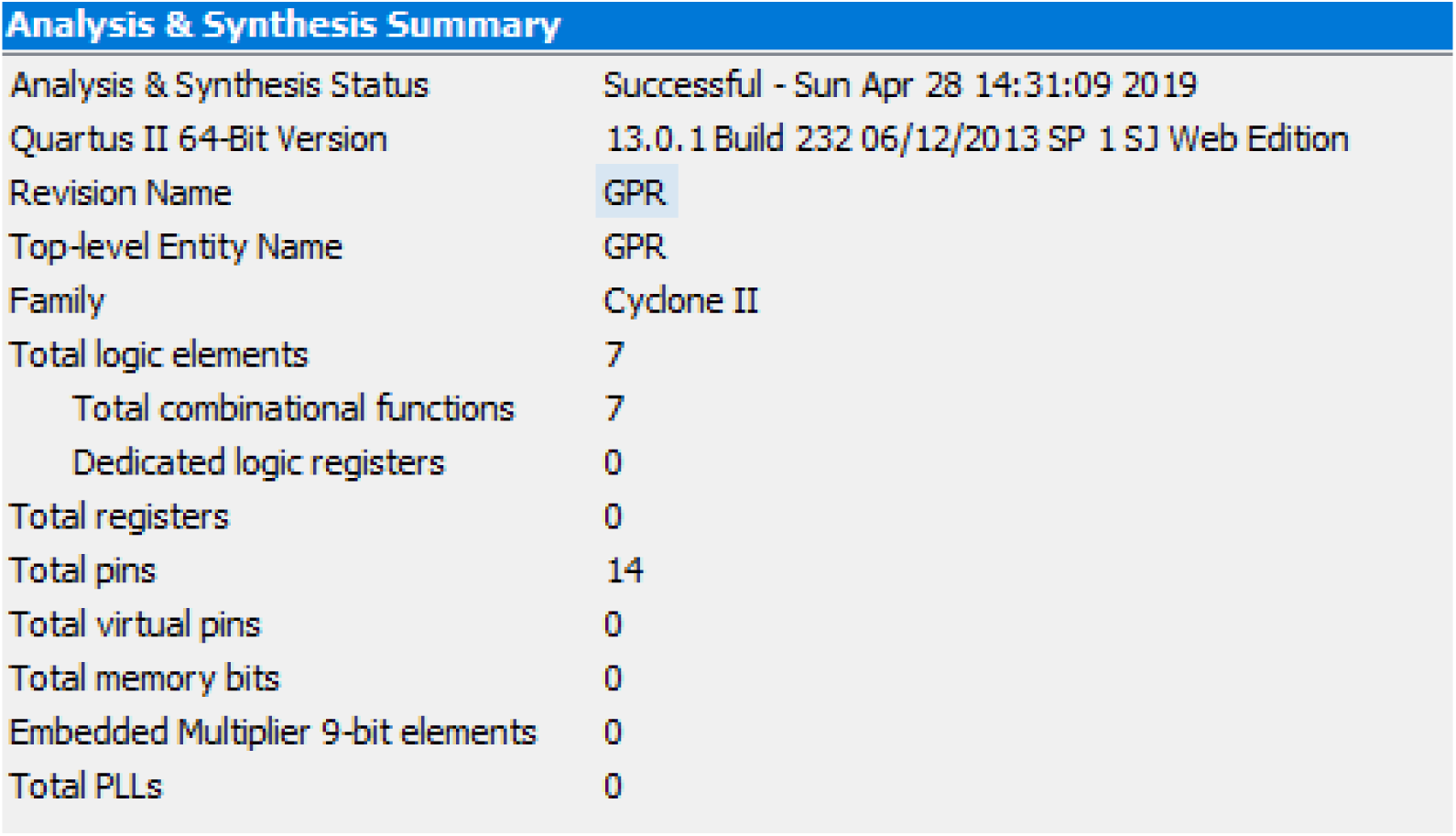
Analysis and Synthesis Summary.

## 5 Discussion

From the results it can be seen that modelling GPRAs on an FPGA could be a very promising first step in creating genome scale metabolic models on hardware. This is given that modelling GPRAs is very natural on an FPGA as they are boolean rules that can be implemented well using in-built logic in FPGA devices. Furthermore it can be seen that FPGAs provide an advantage in terms of speed compared to software implemented models given that the hardware in the FPGA will be fully dedicated to the model. From Figure 2 it is seen that all combinations of gene knock-in and knock-out were completed in 989 ns. However for a fully fledged genome scale model to be implemented on FPGAs more research will have to be done to implement Flux Balance Analysis on these devices which we will investigate in future studies.

## 6 Supplementary Materials

### VHDL Code can be found in GitHub

https://github.com/mcoggins96/Modelling-Gene-Protein-Reaction-Associations-on-an-FPGA

## 7 Dedication

This paper is dedicated to Francesca Marcovecchio who supported me during the writing of this paper.

## Notes

#### Summary of Updates

Figure 2 corrected as original figure contained a mistake regarding LDH_lm reaction that used the GPR (HGNC6541 and HGNC6535) or HGNC19708 instead of the correct GPR HGNC6541 or HGNC6535 or HGNC19708

https://github.com/mcoggins96/Modelling-Gene-Protein-Reaction-Associations-on-an-FPGA

